# Developmental plasticity of *Brachypodium distachyon* in response to P deficiency: modulation by inoculation with phosphate-solubilizing bacteria

**DOI:** 10.1101/858233

**Authors:** Caroline Baudson, Benjamin M. Delory, Stijn Spaepen, Patrick du Jardin, Pierre Delaplace

**Affiliations:** Plant Sciences, Gembloux Agro-Bio Tech, University of Liège, Belgium; Institute of Ecology, Leuphana University, Lüneburg, Germany; Leuven Institute for Beer Research, University of Leuven, Belgium

**Keywords:** biomass allocation, root system morphology, P use efficiency, bio-inoculants, P solubilizing bacteria

## Abstract

**Background:** Mineral P fertilisers must be used wisely in order to preserve rock phosphate, a limited and non-renewable resource. The use of bio-inoculants to improve soil nutrient availability and trigger an efficient plant response to nutrient deficiency is one potential strategy in the attempt to decrease P inputs in agriculture.

**Method:** A gnotobiotic co-cultivation system was used to study the response of *Brachypodium distachyon* to contrasted P supplies (soluble and poorly soluble forms of P) and inoculation with P solubilizing bacteria. *Brachypodium*’s responses to P conditions and inoculation with bacteria were studied in terms of developmental plasticity and P use efficiency.

**Results:** *Brachypodium* showed plasticity in its biomass allocation pattern in response to variable P conditions, specifically by prioritizing root development over shoot productivity under poorly soluble P conditions. Despite the ability of the bacteria to solubilize P, shoot productivity was depressed in plants inoculated with bacteria, although the root system development was maintained. The negative impact of bacteria on biomass production in *Brachypodium* might be attributed to inadequate C supply to bacteria, an increased competition for P between both organisms under P-limiting conditions, or an accumulation of toxic bacterial metabolites in our cultivation system. Both P and inoculation treatments impacted root system morphology. The modulation of *Brachypodium*’s developmental response to P supplies by P solubilizing bacteria did not lead to improved P use efficiency.

**Conclusion:** Our results support the hypothesis that plastic responses of *Brachypodium* cultivated under P-limited conditions are modulated by P solubilizing bacteria. The considered experimental context impacts plant–bacteria interactions. Choosing experimental conditions as close as possible to real ones is important in the selection of P solubilizing bacteria. Both persistent homology and allometric analyses proved to be useful tools that should be considered when studying the impact of bio-inoculants on plant development in response to varying nutritional context.

## 1 Introduction

An important challenge for this century is to implement sustainable cropping systems that preserve the environment and non-renewable resources. This is particularly true concerning phosphate rock, a finite resource mined from only a few countries (Cooper et al. 2011). Phosphate rock constitutes the main source of phosphorus inorganic fertiliser and has been extensively used by farmers in industrialized countries during the past century. Given the growing demand for food, the increasing demand for P fertilisers is predicted to continue (mainly in the developing countries). Although this is a controversial issue, it is estimated that global commercial P reserves could be depleted within 50 to 100 years (Cordell et al. 2009). In the foreseeable future, a peak in P production is expected in 2033. This implies that the growing demand for P will overtake the economically available supply due to decreasing quality and accessibility of the remaining phosphate rock reserves (Cordell and White 2011). As the European Union is strongly dependent on P imports (van Dijk et al. 2016), phosphate rock has been classified as a “critical raw material” by the European Commission since 2014 (European Commission 2014). These considerations emphasize the necessity of adapting fertilisation strategies.

In industrialized countries, the excessive use of P fertilisers has led to an accumulation of P in agricultural soils, constituting a new source of P reserves known as “legacy soil P” (van Dijk et al. 2016; Menezes-Blackburn et al. 2018; Rowe et al. 2016). In some regions, this accumulation has reached levels that generate an environmental risk of watercourse contamination and subsequent eutrophication (Haygarth et al. 2014). It has been estimated that the total soil P stock for arable and grassland soils represents 352 ± 26 years of agronomic P use, with orthophosphate and monoester (organic) phosphate accounting for the greatest proportion (study based on 258 different soils collected in Europe, Oceania and North America; Menezes-Blackburn et al. 2018). However, this soil P reserve is unavailable to plants due to the capacity of many soils to fix P (Shen et al. 2011). Soil P reserve mobilizing technologies must be developed in order to reduce the use of inorganic P fertilisers. The requirement for inorganic P fertilisers could be reduced by 50% if legacy soil P was included in nutrient management practices (Sattari et al. 2012).

Plants have developed strategies to cope with P deficiency and enhance their P use efficiency (PUE), including alteration of the root morphology and architecture, as well as exudation of carboxylates and hydrolytic enzymes for P solubilization. Micro-organisms can also be useful in mobilizing soil P reserves, by directly increasing the P availability in soils (through solubilization) or enhancing plant P nutrition processes (through hormonal stimulation of root growth, for example) (Richardson et al. 2011). The use of micro-organisms, broadly called biological inoculants (“bio-inoculants”), is considered a technology to improve soil P use by crops and pastures (Owen et al. 2015). Those products belong to the biostimulants category as defined by du Jardin (2015). Biostimulants including substances and/or microorganisms act in addition to fertilisers, with the aim of optimising the efficiency of those fertilisers and reducing nutrient application rates (European Parliament and Council of the EU 2019).

Plants exhibit modular growth, potentially allowing them to add new branches to their body. New meristems are exposed to the direct influence of the environment, making plant growth a flexible process (Schmid 1992). Phenotypic plasticity, i.e. the environment-driven alteration of a phenotype, gives the plant a great potential to respond to fluctuating environments (Nicotra et al. 2010; Schmid 1992). It is, at least in part, genetically controlled and heritable (Nicotra et al. 2010). Breeding programs have traditionally opted for phenotypic stability over plasticity, ensuring high yield in constant agricultural systems with high inputs. However, the uncertainty of the future environment and climate requires us to reconsider the place of phenotypic plasticity in breeding strategies. Indeed, phenotypic plasticity can be an advantage to plants living in changing or heterogeneous environments by increasing plant fitness (Lobet et al. 2019; Nicotra et al. 2010). The plastic response of plants to abiotic factors and to the presence of microorganisms still needs clarification. Additionally, the microbial-triggered change in plants’ plastic response to P deficiency deserves greater attention in order to optimize plant P nutrition and reduce the use of fertilisers (Goh et al. 2013).

This study aimed to characterize *Brachypodium distachyon*’s response to contrasted P supplies (soluble and poorly soluble forms of P), as well as the impact of plant inoculation with single strains of phosphate-solubilizing bacteria (PSB) on this response in terms of developmental plasticity. The following hypotheses were tested: (i) biomass allocation and root system development in *Brachypodium* show plasticity in response to contrasted P conditions; (ii) inoculation with PSB modulates the plant’s plastic response to contrasted P supplies; and (iii) this modulation induces changes in plant PUE. Biomass accumulation and allocation, shoot P concentration and PUE, as well as root architectural traits, were considered. *Brachypodium*’s developmental plasticity was assessed using tools including allometry analysis for the biomass allocation and persistent homology analysis for the root system architecture (RSA). It is the first time, to our knowledge, that these tools have been used to precisely evaluate the impact of biostimulants on a plant’s response to nutrient limitation.

## 2 Material and Methods

### 2.1 Plant and bacterial material

*Brachypodium distachyon* (L.) P. Beauv. (Bd21 line) caryopses were kindly provided by Dr Philippe Vain from the John Innes Centre (Norwich, UK) and propagated under greenhouse conditions.

Four bacterial strains were selected for their potential plant growth promotion and phosphorus solubilization capacities: *Bacillus velezensis* GB03 (BveGB03), *Bacillus velezensis* FZB42 (BveFZB42), *Pseudomonas fluorescens* 29ARP (Pfl29ARP) and *Azotobacter vinelandii* F0819 (AviF0819). The *Bacillus* strains were selected for their plant growth promotion activities on *Poaceae* (Delaplace et al. 2015; Myresiotis et al. 2015; Zhang et al. 2014), as well as their ability to solubilize different forms of P (Giles et al. 2014; Idris et al. 2007; Idriss et al. 2002; Liu et al. 2015). *Pseudomonas fluorescens* also exhibited P solubilizing activities and promoted wheat (Shaharoona et al. 2008) and maize growth (Li et al. 2017). *Azotobacter vinelandii*, a free diazotrophic bacteria, exhibited P solubilization activity (Nosrati et al. 2014) and PGP traits (Taller and Wong 1989). *Escherichia coli* DH5α 99B829 (Eco99B829), was selected as a negative control for plant growth promotion (Delaplace et al. 2015; Wu et al. 2016; Zhou et al. 2016). The strains BveGB03 and Eco99B829 were kindly provided by Dr Paul W. Paré and Dr John McInroy (Texas Tech University, Lubbock, TX, USA), Pfl29ARP by Dr Alain Sarniguet (Institut National de la Recherche Agronomique, Rennes, France), AviF0819 by the Katholieke Universiteit Leuven (Leuven, Belgium), and BveFZB42 by Pr Rainer Borriss (Nord Reet UG, Greifswald, Germany). The bacterial strains were stored at 80°C in LB medium containing 20% v/v glycerol before plating.

### 2.2 *In vitro* P solubilization assay

One week before the experiment, the bacteria were plated on LB agar plates (2.5% w/v LB broth, Prod. No. L3152; 1.5% w/v agar, Prod. No. 05039, Sigma-Aldrich Co., St. Louis, USA) and incubated at 28°C. The day before the experiment, the bacteria were suspended in 40 ml of LB (2.5% w/v LB broth) and incubated overnight at 150 rpm and 30°C (Innova 4340, New Brunswick Scientific Co. Inc., Edison, USA). The concentration of the bacterial suspensions was derived from the optical density, measured at 540 nm. The tubes were centrifuged (20 min at 4000 rpm) and the LB medium was removed. The bacterial pellets were rinsed with 25 ml of 10 mM MgSO_4_ in order to avoid P contaminations. The tubes were centrifuged again (20 min at 4000 rpm) and the MgSO_4_ solution was removed. The bacteria were suspended in an adequate volume of NBRIP medium (National Botanical Research Institute’s phosphate growth medium, Nautiyal 1999) containing tricalcium phosphate (Ca_3_(PO_4_)_2_, hereafter named “TCP”, Prod. No. C0506.1000, Duchefa Biochemie, Haarlem, The Netherlands) or hydroxyapatite (Ca_5_(PO_4_)_3_OH, hereafter named “HA”, Prod. No. 8450.1, Carl Roth GmbH + Co. KG, Karlsruhe, Germany) at a concentration of 5 g/l (pH 7) in order to obtain a bacterial concentration of 10^7^ CFU/ml. Bottles containing 90 ml of NBRIP medium were successively inoculated with 10 ml of the prepared suspensions to obtain a final concentration of 10^6^ CFU/ml, and incubated for 3 days at 30°C and 150 rpm.

10 ml were sampled daily for subsequent analysis. 1 ml was subsampled for serial dilution and plating on LB agar plates in order to monitor bacterial growth. The remaining samples were centrifuged and the supernatant was filter-sterilized (pore size 0.2 μm) for pH and soluble P content measurements. The P content in the solution (as soluble phosphate) was measured according to the phosphomolybdate blue colorimetric method (Murphy and Riley 1962) (Prod. No. 69888, molybdate reagent solution, Fluka Sigma-Aldrich Co., St. Louis, USA).

### 2.3 *Brachypodium*-bacteria co-cultivation in axenic conditions

One week before the experiment, the bacteria were plated on LB agar plates and incubated at 28°C. The day before the experiment, the bacteria were suspended in 40 ml of LB and incubated overnight at 150 rpm and 30°C. The tubes were centrifuged (20 min at 4000 rpm) and the LB medium was removed. Inoculums at 10^8^ CFU/ml were finally prepared in 10 mM MgSO_4_ for subsequent inoculation of the plantlets.

*Brachypodium distachyon* Bd21 seeds were surface sterilized (30 s in 70% v/v ethanol, rinsed once with sterile water, 10 min in sodium hypochlorite 5% v/v, rinsed three times with sterile water) and stratified for 2 days at 4°C on Hoagland agar plates (0.125% w/v Hoagland, Prod. No. DU1201, Duchefa Biochemie, Haarlem, The Netherlands; 0.094% w/v Ca(NO_3_)_2_.4H_2_O; 0.8% w/v Plant agar, Prod. No. P1001, Duchefa Biochemie, Haarlem, The Netherlands). The seeds were then incubated for 24 hours in a growth chamber (23°C, 16h/8h day light, PPFD 140 μmol.m^−2^.s^−1^) for germination.

Homogeneous 24 hour-old plantlets were selected and inoculated with bacteria by dipping them into 10 mM MgSO_4_ containing a bacterial strain at 10^8^ CFU/ml for 10 minutes (control plantlets were dipped into 10 mM MgSO_4_). The plantlets were then transferred into Magenta^®^ boxes (GA-7 Magenta vessel, Magenta LLC, Lockport, USA) filled with 180 g of sterilized black gravel (rinsed three times with tap water and autoclaved; 1-3 mm quartz gravel, prod. no. 400723, Flamingo, Geel, Belgium) and 50 ml of sterile nutrient solution. One plantlet was placed into each Magenta^®^ box. Three modified Hoagland nutrient solutions and a reference solution, corresponding to the contrasting P treatments, were used: a P-limiting supply containing 25μM of KH_2_PO_4_ (“P-”), a P-limiting supply supplemented with 1 g/l TCP (“P-/TCP”) or 1 g/l HA (“P-/HA”), and a P-sufficient supply containing 1mM KH_2_PO_4_ (“P+”). The boxes were sealed with Leukopor^®^ tape (prod.no. 02454-00, BSN medical GmbH, Hamburg, Germany) and incubated in the growth chamber for four weeks (23°C, 16h/8h day light, PPFD 140 μmol.m^−2^.s^−1^). Six independent experiments were performed (three with P-, P-/TCP and P+ treatments; three with P-, P-/HA and P+ treatments) and five plants were cultivated for each treatment (90 plants per experiment). Four week-old plants were harvested and cut to measure fresh biomass accumulation in shoot and roots. The presence of bacteria was assessed by scratching agar plates with the root system. The root system was scanned for three plants per treatment (1200 dpi, Epson Perfection V800 Photo, Epson America Inc., Long Beach, USA) in order to perform RSA analyses. Shoots were stored at −80°C before P content measurements. Total biomass and root mass fraction (hereafter named “RMF”, mg root biomass/mg total biomass) were computed from the measured biomasses. RMF was recorded in order to analyse biomass allocation in *Brachypodium*, considering allocation as a partitioning process. According to this perspective, plants divide a given amount of resources among structures according to their developmental priorities (Weiner 2004).

### 2.4 P concentration in plant tissues

P content in *Brachypodium* shoots was measured by ICP-OES on frozen samples (C.A.R.A.H. ASBL, Ath, Belgium). The samples were calcinated overnight at 450°C. The ashes were then suspended in nitric acid for digestion. The P concentration was measured by ICP-OES (Thermo Fisher iCAP 7600, Thermo Fisher Scientific, Waltham, USA). The five replicates of each treatment in each independent experiment were pooled. Three pooled samples were analysed for the P-/TCP and P-/HA treatments. Six pooled samples were analysed for the P- and P+ treatments. The results were expressed as total shoot P concentration (μg P/ mg fresh weight).

### 2.5 Root system architecture measurement

An automated evaluation of the total root length (“TRL”) was performed for all scanned root systems using the ImageJ macro IJ_Rhizo (Pierret et al. 2013; Schneider et al. 2012). For each image, the TRL was estimated using the Kimura method as it provides more accurate length estimates than the other methods available in IJ_Rhizo (Delory et al. 2017).

In addition, more detailed root system architecture analyses were performed using SmartRoot (Lobet et al. 2011). Only the 1^st^ and 2^nd^ order roots were analysed because the thinner, higher order, roots break easily at harvest. These manual analyses were performed for the control treatment and the two strains found to impact plant development the most. The RSML (Root System Markup Language) outputs were then processed with the archiDART package for morphological analysis using persistent homology (R 3.5.2, R core Team 2018; archiDART package version 3.3, Delory *et al*., 2016; Delory et al. 2018). A geodesic distance function was used to compute a persistence barcode for each root system. The degree of dissimilarity between barcodes (i.e. root systems) was assessed by computing a pairwise distance matrix containing dissimilarities calculated using a bottleneck distance method. Morphological differences between root systems were then visualized using multidimensional scaling (R 3.5.2, R Core Team 2018).

### 2.6 P use efficiency

The P use efficiency analysis was performed by considering three different parameters: (i) the P uptake efficiency (PUpE, μg P/mg P applied), corresponding to the shoot P content per unit of soluble P applied; (ii) the P utilization efficiency (PUtE, mg FW/μg P), corresponding to the biomass produced by unit of shoot P; and (iii) the physiological P use efficiency (PPUE, mg^2^ FW/μg P), corresponding to the produced biomass divided by the tissue P concentration (Neto et al. 2016).

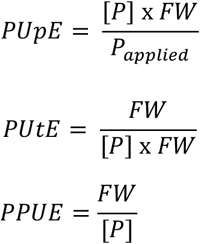

### 2.7 Statistical analyses

The relationship between P solubilization and pH variation in the NBRIP medium was studied by performing regression analyses (lm function, R 3.5.2, R Core Team 2018). The model order was increased until there was not significant difference with the higher order model (anova function, R 3.5.2, R Core Team 2018).

Three-way ANOVAs were performed to study the impact of P supply, bacteria inoculation, independent temporal repetitions and the interaction between P supply and bacteria inoculation on the following factors: shoot, root and total biomass parameters; RMF; TRL; shoot P concentration; and the three components of PUE. A model with crossed fixed factors was applied (lm, glm and anova functions, R 3.5.2, R Core Team 2018; Gamma family distribution with a log-link function was used for GLM models). Dunnett’s post-hoc tests were performed to compare the treatments to the control situation (non-inoculated plants for inoculation treatment, P+ for P treatment) (R 3.5.2, R Core Team 2018; multcomp package version 1.4-8, Hothorn et al. 2008).

Allometry analyses were performed on shoot and root biomass in order to study the biomass allocation pattern. The “smatr” package (R 3.5.2, “smatr” version 3.4-8) was used for estimation, inference and plotting of allometric lines as well as for checking assumptions (Warton et al. 2012). The standardized major axis (“SMA”) analysis was used and all variables were log-transformed. In brief, this analysis consists of a model II regression, estimating how one variable scales against another. The obtained allometric trajectories depict the relative development of the shoot and root compartments, i.e. how the root system growth impacts the shoot development. Inference statistics compare coefficients of the regression lines (slope and elevation) between the populations (Warton et al. 2006). Firstly, differences in slope between groups were tested. If there was no difference in slope between groups, differences in elevation were tested using a common slope for all groups. When significant differences between groups were highlighted, pairwise multiple comparisons were performed in order to identify which populations differed from each other. Differences in slope (i.e. investment in shoot biomass per additional unit of root biomass) or elevation (i.e. shoot productivity for similar root biomass) among treatments led to different allometric trajectories. Change in allometric trajectory due to different treatments revealed plasticity in the biomass allocation process (Weiner 2004; Xie et al. 2015). The analysis of allometric trajectories is complementary to the analysis of RMF for the study of biomass allocation plasticity.

Differences in root system architecture were investigated using permutational multivariate analysis of variance (PERMANOVA) (R 3.5.2, R Core Team 2018; vegan package version 2.5-4, Oksanen et al. 2019). The dissimilarity matrix used in the model formula was the pairwise distance matrix returned by the persistent homology analysis of plant root systems. Bacterial strain, P treatment and their interaction were used as independent variables in the model. For each fixed factor, a post-hoc test was performed by running a separate PERMANOVA for each pairwise comparison. *P* values were adjusted for multiple comparisons using the Bonferroni method.

All figures shown in this study were generated using the “ggplot2” package (R 3.5.2, “ggplot2” version 3.1.0).

Data and R scripts are accessible on Zenodo repository at https://doi.org/10.5281/zenodo.3555566

## 3 Results

### 3.1 The selected bacterial strains solubilized tricalcium phosphate and hydroxyapatite, while acidifying their growth medium

The bacteria’s ability to solubilize poorly available forms of P was assessed using TCP and HA in a modified NBRIP medium. After three days, all the selected strains were able to solubilize both forms of P to some extent (Fig. 1a) compared to the non-inoculated control treatment. For both forms of P, the best performing bacterial strains were Eco99B829 and Pfl29ARP. The solubilization of TCP and HA were similar for all bacterial strains with the exception of Eco99B829, which exhibited a stronger solubilization ability for HA despite a greater variability between independent replicates. All the strains were able to maintain stationary populations during the duration of the experiment (data not shown). BveGB03, AviF0819, Pfl29ARP and Eco99B829 generated a pH drop during the experiment for both forms of P (Fig. 1a). As for the P concentration, Eco99B829 and Pfl29ARP induced the strongest acidification.

**Fig. 1.**
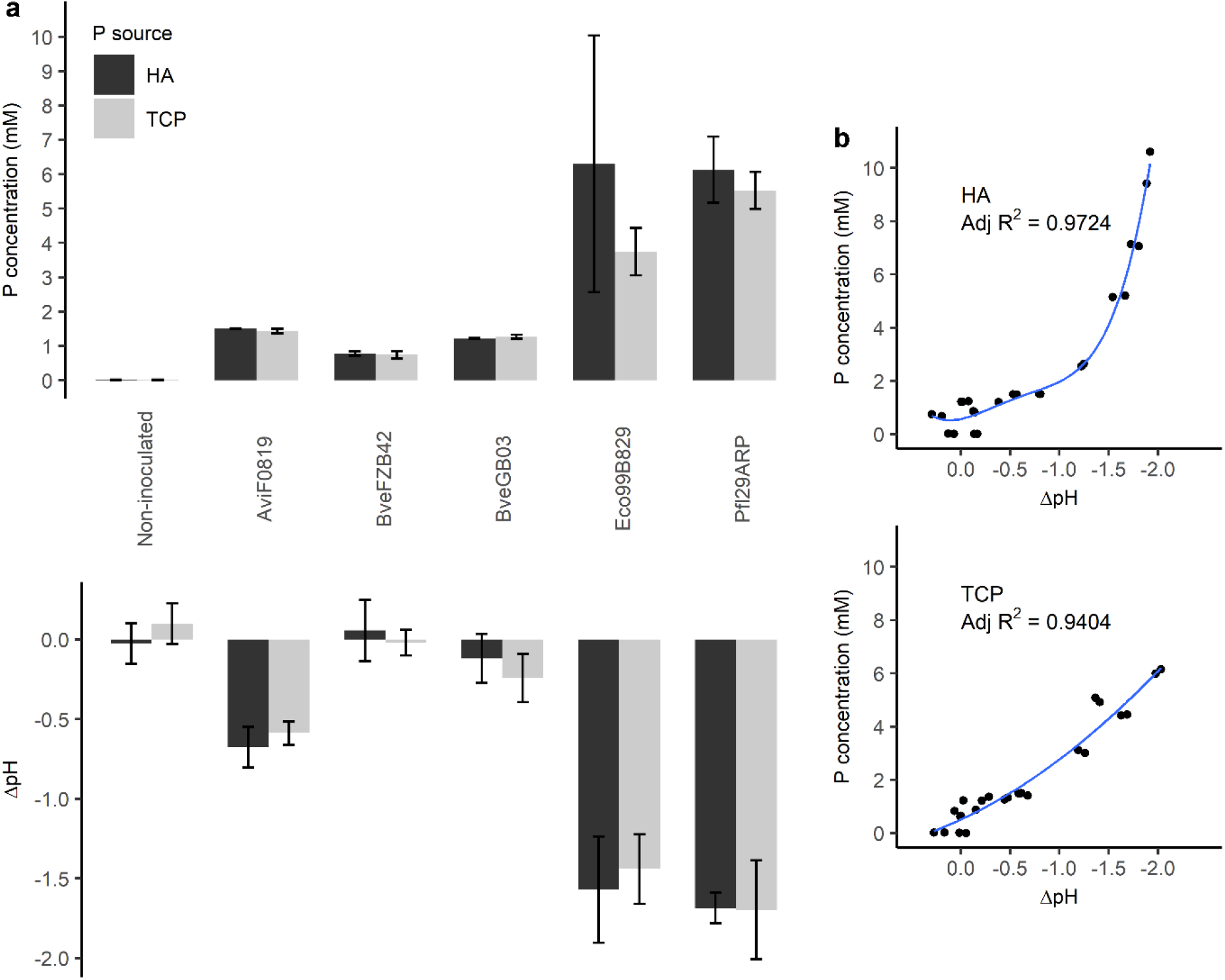
(a) Soluble P concentration and pH variation in NBRIP medium after three days of bacteria cultivation in the presence of either TCP or HA as poorly soluble forms of P (n=4, mean ± SD). (b) Regression curves linking the observed P concentration and the ΔpH in the growing medium after three days of incubation. For each regression model, adjusted R^2^ values are displayed on the graphs. Regression coefficients for HA solubilization: y = 0.571 – 0.814x + 2.7675x^2^ + 4.219x^3^ + 2.039x^4^; regression coefficients for TCP solubilization: y = 0.513 – 1.727x + 0.530x^2^

Regarding HA solubilization, the relationship between the soluble P concentration and ΔpH in the growing medium was best fitted by a 4^th^ order polynomial model (Fig. 1b). The HA solubilization activity clearly intensified as the acidification became stronger. The regression between soluble P concentration and ΔpH for TCP solubilization was best fitted by a 2^nd^ order polynomial model (Fig. 1b). As for HA, the TCP solubilization activity intensified with increasing pH variation, but to a lesser extent.

### 3.2 Biomass accumulation in *Brachypodium* was altered by soluble P deficiency and inoculation with P solubilizing bacteria

Shoot biomass production was lower in plants grown under P-, P-/TCP and P-/HA conditions, with a diminution of 43.1%, 35.2% and 33.4% compared to the P+ treatment, respectively (*P*<0.001, Fig. 2a-d). Plant inoculation with PSB strains had either no impact or induced a lower shoot biomass accumulation (Fig. 2e). Inoculation with BveFZB42 and Pfl29ARP led to a significantly lower shoot biomass, with up to 13.2% reduction in plants inoculated with BveFZB42 under the P+ treatment and 30.3% reduction in plants inoculated with Pfl29ARP under the P-/TCP treatment (*P*<0.001). The impact of P conditions on the accumulation of biomass in roots was more limited, with only plants grown under the P- treatment having a significantly greater root biomass (+13.3%) compared to the plants exposed to P+ conditions (*P*<0.001, Fig. 2f-i). Inoculation had either no impact or a negative impact on the accumulation of biomass in roots (*P*=0.003, Fig. 2j). Indeed, plants inoculated with BveFZB42 exhibited a significant reduction of the root biomass of up to 14.5% under the P- treatment. The total biomass decreased by 27.8%, 25.2% and 24.1% under the P-, P-/TCP and P-/HA treatments respectively (*P*<0.001 Fig. 2k-n). Plant inoculation with PSB strains led to a repression of the biomass accumulation at the whole plant level (*P*<0.001, Fig. 2o). Inoculation with BveFZB42 and Pfl29ARP induced a significantly lower total biomass in comparison with the non-inoculated control. The growth reduction reached 12.2% with BveFZB42 under P-/HA conditions and 21.1% with Pfl29ARP under P- conditions. A table comprising mean values per treatment, standard deviation and coefficients of ANOVAs is available in Online Resource 1.

**Fig. 2.**
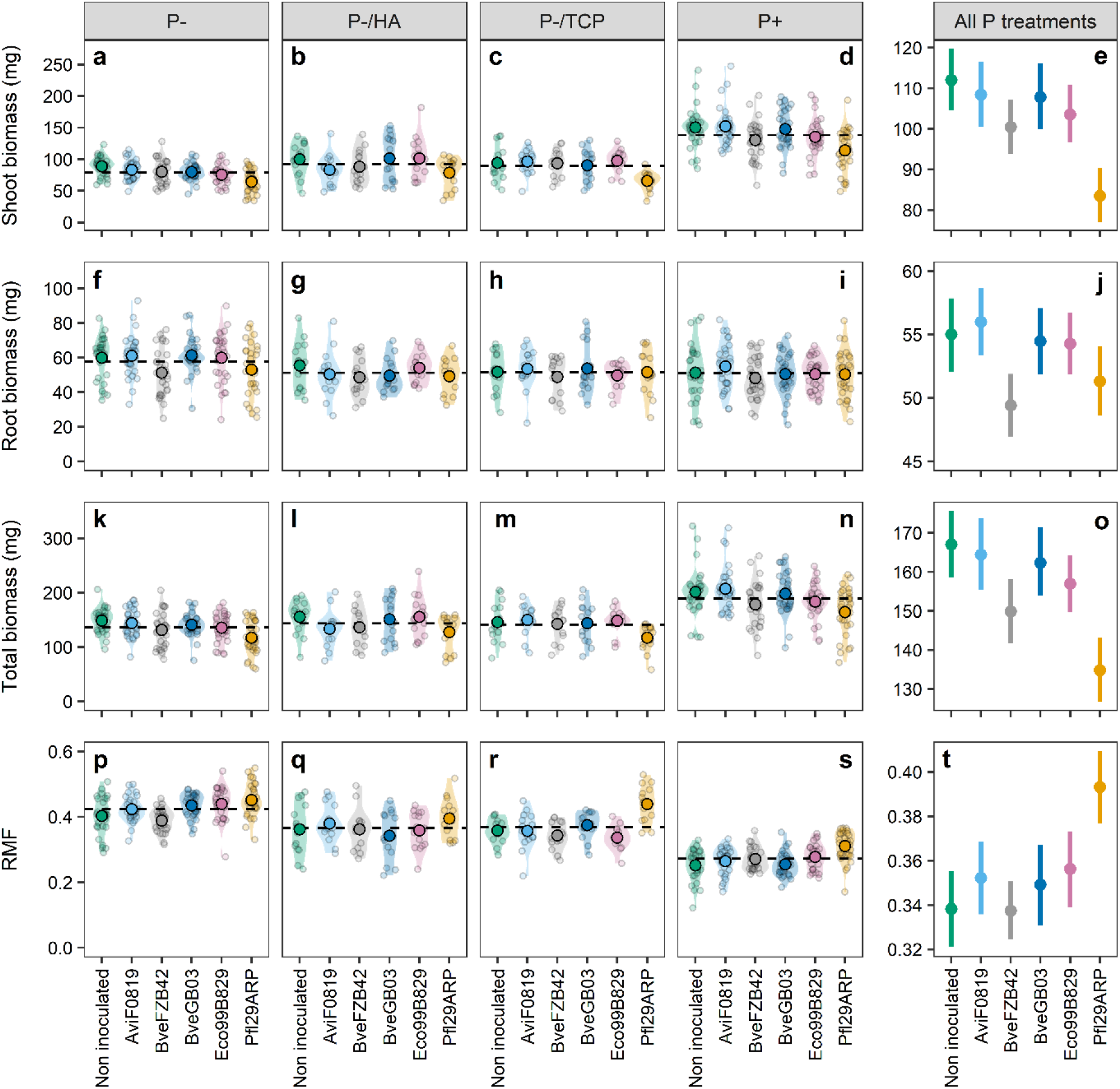
Average shoot biomass (a-e), root biomass (f-j), total biomass (k-o) and root mass fraction (p-t) of four-week-old *Brachypodium* plantlets exposed to contrasted P supplies and either inoculated or not inoculated with bacteria. n = 30 for the P- and P+ treatments, n=15 for the P-/HA and P-/TCP treatments. For each P treatment, the grand mean is shown by a dashed horizontal line. For each inoculation treatment, large black-circled dots represent mean values, and shaded areas show the density distribution of each population. Individuals are displayed as small grey-circled dots in the graphs. In panels e, j, o and t, values are means +/- 95% confidence intervals calculated across P treatments

**Online Resource 1.**
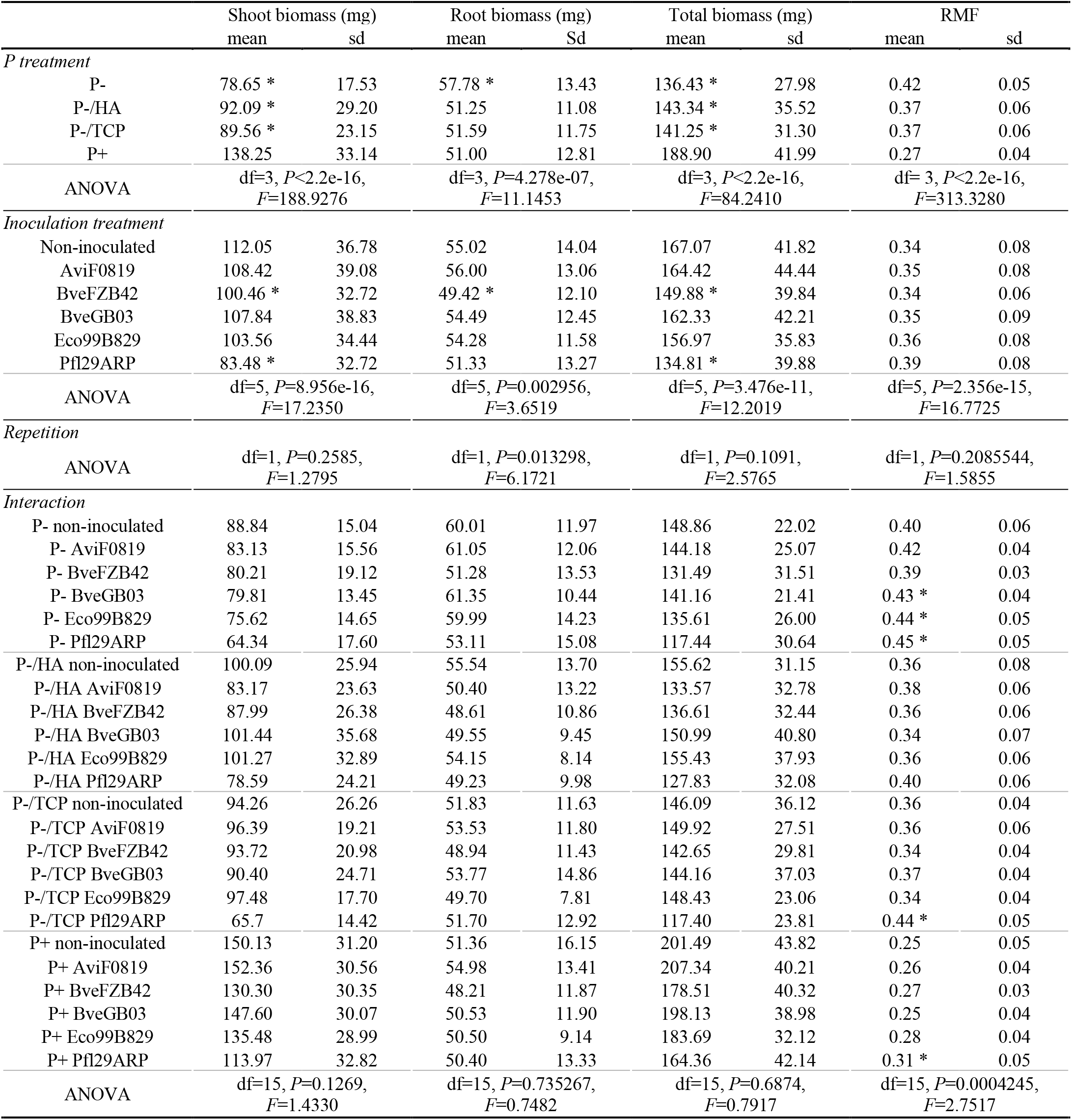
Biomass accumulation and RMF of four-week-old *Brachypodium* plantlets grown in Magenta boxes, exposed to contrasted P supplies and either inoculated or not inoculated with bacterial strains (n=30 for the P- and P+ treatments, n=15 for the P-/TCP and P-/HA treatments). Results of 3-way ANOVAs (degree of freedom “df”, *P* and *F* values) and Dunnett’s *post hoc* tests (annotated with stars; P+ and non-inoculated treatments used as references)

|  | Shoot biomass (mg) |  | Root biomass (mg) |  | Total biomass (mg) |  | RMF |  |
| --- | --- | --- | --- | --- | --- | --- | --- | --- |
|  | mean | sd | mean | Sd | mean | sd | mean | sd |
| *P treatment* |  |  |  |  |  |  |  |  |
| P- | 78.65 * | 17.53 | 57.78 * | 13.43 | 136.43 * | 27.98 | 0.42 | 0.05 |
| P-/HA | 92.09 * | 29.20 | 51.25 | 11.08 | 143.34 * | 35.52 | 0.37 | 0.06 |
| P-/TCP | 89.56 * | 23.15 | 51.59 | 11.75 | 141.25 * | 31.30 | 0.37 | 0.06 |
| P+ | 138.25 | 33.14 | 51.00 | 12.81 | 188.90 | 41.99 | 0.27 | 0.04 |
| ANOVA | df=3, *P*<2.2e-16,<br>*F*=188.9276 |  | df=3, *P*=4.278e-07,<br>*F*=11.1453 |  | df=3, *P*<2.2e-16,<br>*F*=84.2410 |  | df= 3, *P*<2.2e-16,<br>*F*=313.3280 |  |
| *Inoculation treatment* |  |  |  |  |  |  |  |  |
| Non-inoculated | 112.05 | 36.78 | 55.02 | 14.04 | 167.07 | 41.82 | 0.34 | 0.08 |
| AviF0819 | 108.42 | 39.08 | 56.00 | 13.06 | 164.42 | 44.44 | 0.35 | 0.08 |
| BveFZB42 | 100.46 * | 32.72 | 49.42 * | 12.10 | 149.88 * | 39.84 | 0.34 | 0.06 |
| BveGB03 | 107.84 | 38.83 | 54.49 | 12.45 | 162.33 | 42.21 | 0.35 | 0.09 |
| Eco99B829 | 103.56 | 34.44 | 54.28 | 11.58 | 156.97 | 35.83 | 0.36 | 0.08 |
| Pfl29ARP | 83.48 * | 32.72 | 51.33 | 13.27 | 134.81 * | 39.88 | 0.39 | 0.08 |
| ANOVA | df=5, *P*=8.956e-16,<br>*F*=17.2350 |  | df=5, *P*=0.002956,<br>*F*=3.6519 |  | df=5, *P*=3.476e-11,<br>*F*=12.2019 |  | df=5, *P*=2.356e-15,<br>*F*=16.7725 |  |
| *Repetition* |  |  |  |  |  |  |  |  |
| ANOVA | df=1, *P*=0.2585,<br>*F*=1.2795 |  | df=1, *P*=0.013298,<br>*F*=6.1721 |  | df=1, *P*=0.1091,<br>*F*=2.5765 |  | df=1, *P*=0.2085544,<br>*F*=1.5855 |  |
| *Interaction* |  |  |  |  |  |  |  |  |
| P- non-inoculated | 88.84 | 15.04 | 60.01 | 11.97 | 148.86 | 22.02 | 0.40 | 0.06 |
| P- AviF0819 | 83.13 | 15.56 | 61.05 | 12.06 | 144.18 | 25.07 | 0.42 | 0.04 |
| P- BveFZB42 | 80.21 | 19.12 | 51.28 | 13.53 | 131.49 | 31.51 | 0.39 | 0.03 |
| P- BveGB03 | 79.81 | 13.45 | 61.35 | 10.44 | 141.16 | 21.41 | 0.43 * | 0.04 |
| P- Eco99B829 | 75.62 | 14.65 | 59.99 | 14.23 | 135.61 | 26.00 | 0.44 * | 0.05 |
| P- Pfl29ARP | 64.34 | 17.60 | 53.11 | 15.08 | 117.44 | 30.64 | 0.45 * | 0.05 |
| P-/HA non-inoculated | 100.09 | 25.94 | 55.54 | 13.70 | 155.62 | 31.15 | 0.36 | 0.08 |
| P-/HA AviF0819 | 83.17 | 23.63 | 50.40 | 13.22 | 133.57 | 32.78 | 0.38 | 0.06 |
| P-/HA BveFZB42 | 87.99 | 26.38 | 48.61 | 10.86 | 136.61 | 32.44 | 0.36 | 0.06 |
| P-/HA BveGB03 | 101.44 | 35.68 | 49.55 | 9.45 | 150.99 | 40.80 | 0.34 | 0.07 |
| P-/HA Eco99B829 | 101.27 | 32.89 | 54.15 | 8.14 | 155.43 | 37.93 | 0.36 | 0.06 |
| P-/HA Pfl29ARP | 78.59 | 24.21 | 49.23 | 9.98 | 127.83 | 32.08 | 0.40 | 0.06 |
| P-/TCP non-inoculated | 94.26 | 26.26 | 51.83 | 11.63 | 146.09 | 36.12 | 0.36 | 0.04 |
| P-/TCP AviF0819 | 96.39 | 19.21 | 53.53 | 11.80 | 149.92 | 27.51 | 0.36 | 0.06 |
| P-/TCP BveFZB42 | 93.72 | 20.98 | 48.94 | 11.43 | 142.65 | 29.81 | 0.34 | 0.04 |
| P-/TCP BveGB03 | 90.40 | 24.71 | 53.77 | 14.86 | 144.16 | 37.03 | 0.37 | 0.04 |
| P-/TCP Eco99B829 | 97.48 | 17.70 | 49.70 | 7.81 | 148.43 | 23.06 | 0.34 | 0.04 |
| P-/TCP Pfl29ARP | 65.7 | 14.42 | 51.70 | 12.92 | 117.40 | 23.81 | 0.44 * | 0.05 |
| P+ non-inoculated | 150.13 | 31.20 | 51.36 | 16.15 | 201.49 | 43.82 | 0.25 | 0.05 |
| P+ AviF0819 | 152.36 | 30.56 | 54.98 | 13.41 | 207.34 | 40.21 | 0.26 | 0.04 |
| P+ BveFZB42 | 130.30 | 30.35 | 48.21 | 11.87 | 178.51 | 40.32 | 0.27 | 0.03 |
| P+ BveGB03 | 147.60 | 30.07 | 50.53 | 11.90 | 198.13 | 38.98 | 0.25 | 0.04 |
| P+ Eco99B829 | 135.48 | 28.99 | 50.50 | 9.14 | 183.69 | 32.12 | 0.28 | 0.04 |
| P+ Pfl29ARP | 113.97 | 32.82 | 50.40 | 13.33 | 164.36 | 42.14 | 0.31 * | 0.05 |
| ANOVA | df=15, *P*=0.1269,<br>*F*=1.4330 |  | df=15, *P*=0.735267,<br>*F*=0.7482 |  | df=15, *P*=0.6874,<br>*F*=0.7917 |  | df=15, *P*=0.0004245,<br>*F*=2.7517 |  |

### 3.3 Shifts in biomass partitioning and allometric trajectories of *Brachypodium* were observed when exposed to contrasted P supplies and inoculated with P solubilizing bacteria

Exposure of *Brachypodium* to soluble P limitation (P-, P-/HA and P-/TCP) increased RMF by 55.8%, 34.9% and 35.7% compared to the P+ treatment, respectively (*P*<0.001, Fig. 2p-s). The impact of *Brachypodium* inoculation with PSB was dependent on the P environment, as there was a significant interaction between these two variables (*P*<0.001). Plants inoculated with Pf29ARP exhibited the greatest RMF under all treatments (Fig. 2t) and this effect was significant for the P-, P-/TCP and P+ treatments (12.1%, 22.7% and 23.4% increase, respectively). Under P- conditions, plants inoculated with BveGB03 and Eco99B829 also had a significantly greater RMF compared to non-inoculated plants (Fig. 2p-t). Mean values per treatment, standard deviation and coefficients of ANOVAs are available in Online Resource 1.

The allocation pattern between shoots and roots was further analysed using SMA regression models (Fig. 3). In non-inoculated plants grown under P-/TCP and P-/HA conditions, the shoot biomass increase per unit of root biomass was greater than that of non-inoculated plants grown under P- and P+ conditions (slopes: 1.15, 1.19, 0.81 and 0.60, respectively; *P*=0.021; Fig. 3a). Non-inoculated plants grown under P+ conditions exhibited the greatest shoot productivity, but invested the lowest amount of biomass into the shoot per unit of root production. Inoculation of plants grown under P- conditions did not induce a significant difference in slope (*P*=0.757, Fig. 3b). Significant differences in elevation were observed (*P*<0.001), with non-inoculated plants and plants inoculated with BveFZB42 showing the greatest shoot productivity and plants inoculated with Pfl29ARP the lowest, for similar root biomass. Under the P-/HA treatment, slope and elevation did not significantly vary among groups (*P*=0.174 and 0.433, respectively; Fig. 3c), even if plants inoculated with BveGB03 and Eco99B829 showed greater shoot biomass increase per unit of root biomass, when considering the graphical trends. When plants were grown in the presence of TCP, the inoculation with PSB did not significantly affect the slope (*P*=0.835, Fig. 3d). Elevation was significantly altered when plants were inoculated with Pfl29ARP, leading to the lowest shoot productivity for similar root biomass (lowest elevation, *P*<0.001). Significant differences in slope were observed under the P+ treatment (*P*=0.008), with the greatest production of shoot biomass per unit of root production in plants inoculated with Pfl29ARP, Eco99B829 and BveFZB42 (Fig. 3e). SMA coefficients and results of covariance analysis are available in Online Resource 2.

**Fig. 3.**
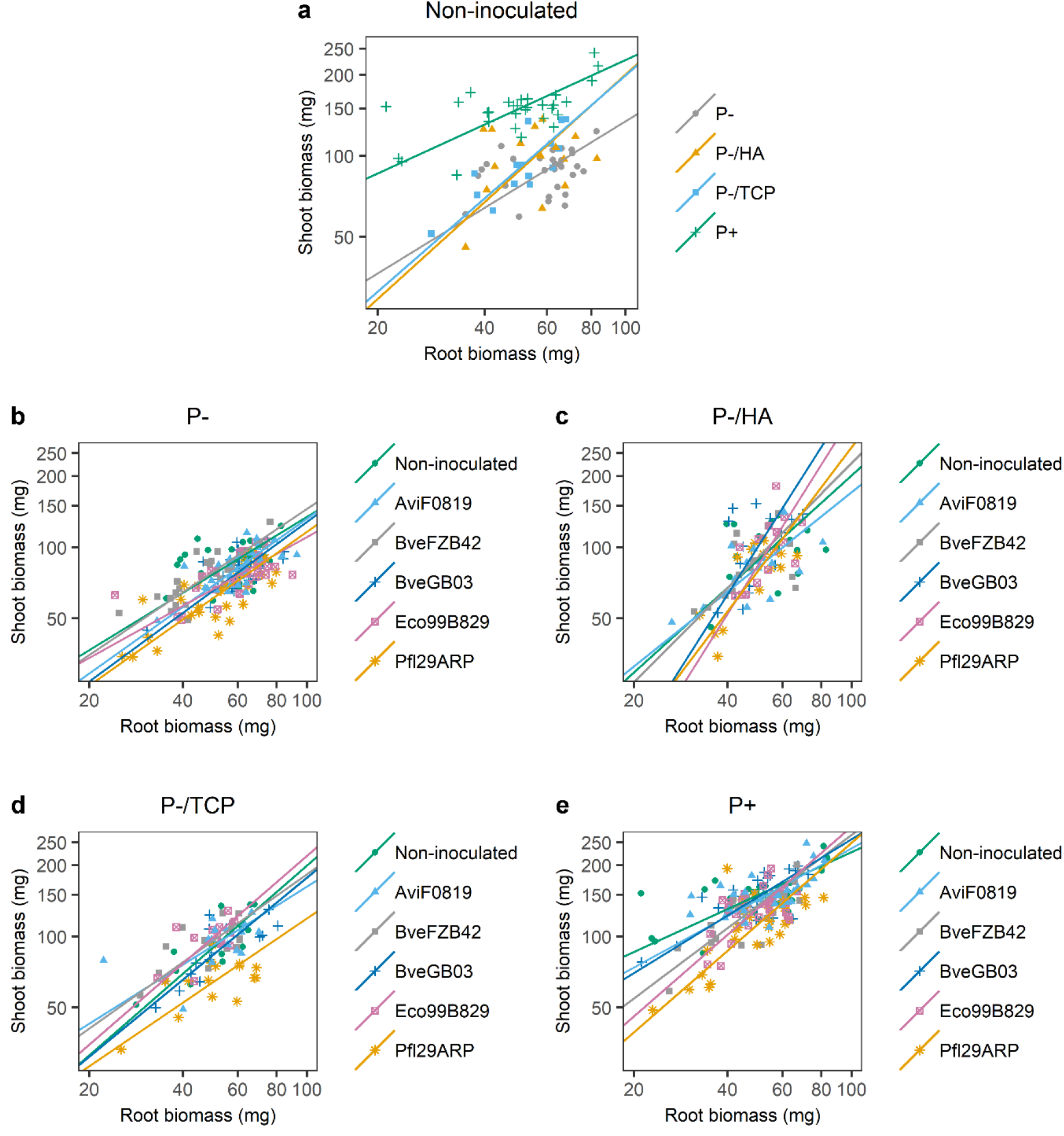
Allometric relationship between shoot biomass and root biomass of four-week-old *Brachypodium* plantlets exposed to contrasted P supplies and grown with or without bacterial inoculation. X and Y axes are log-scaled. Symbols represent individuals. Lines represent SMA regression lines. (a) Non-inoculated plants exposed to contrasted P supplies. Inoculated and non-inoculated plants grown under (b) P-, (c) P-/HA, (d) P-/TCP and (e) P+ conditions

**Online Resource 2.** Coefficients, R^2^ and *P* value of SMA lines (n=30 for the P- and P+ treatments, n=15 for the P-/HA and P-/TCP treatments), results of covariance analysis for differences among SMA lines coefficients (degree of freedom “df”, *P* and likelihood ratio test “*LR*” values). If no significant difference was noticed between slopes, a common slope was used to test for difference in elevation. Treatments without any common letter are significantly different from each other (pairwise comparison)

| | elevation | slope | $R^2$ | $P$ |
| --- | --- | --- | --- | --- |
| *Non-inoculated* |  |  |  |  |
| P- | 0.51 | 0.81 *ab* | 0.090 | 0.10762 |
| P-/HA | -0.08 | 1.19 *a* | 0.072 | 0.33277 |
| P-/TCP | 0.00 | 1.15 *a* | 0.668 | 0.00019 |
| P+ | 1.15 | 0.60 *b* | 0.399 | 0.00018 |
| Covariance analysis | / | df= 3,<br>$P=0.020606$, | | |
| *P-* |  |  |  |  |
| Non-inoculated | 0.38 *ab* |  | 0.090 | 0.10762 |
| AviF0819 | 0.34 *bc* |  | 0.468 | 3.0804e-5 |
| BveFZB42 | 0.39 *a* | 0.88 | 0.718 | 3.5339e-9 |
| BveGB03 | 0.32 *cd* |  | 0.460 | 5.3114e-5 |
| Eco99B829 | 0.31 *cd* |  | 0.411 | 0.00013 |
| Pfl29ARP | 0.28 *d* |  | 0.604 | 4.3258e-7 |
| Covariance analysis | df=5, $P=2.4697e-9$,<br>LR=48.77 | df=5, $P=0.75661$,<br>LR=2.631 | | |
| *P-/HA* |  |  |  |  |
| Non-inoculated | -0.71 |  | 0.072 | 0.33277 |
| AviF0819 | -0.73 |  | 0.394 | 0.01228 |
| BveFZB42 | -0.69 | 1.56 | 0.264 | 0.05028 |
| BveGB03 | -0.65 |  | 0.227 | 0.07242 |
| Eco99B829 | -0.71 |  | 0.384 | 0.01371 |
| Pfl29ARP | -0.75 |  | 0.582 | 0.00093 |
| Covariance analysis | df=5, $P=0.43264$,<br>LR=4.865 | df=5, $P=0.17355$,<br>LR=7.7 | | |
| *P-/TCP* |  |  |  |  |
| Non-inoculated | 0.23 *ab* |  | 0.668 | 0.00019 |
| AviF0819 | 0.23 *ab* |  | 0.249 | 0.05839 |
| BveFZB42 | 0.25 *ab* | 1.02 | 0.513 | 0.00269 |
| BveGB03 | 0.19 *b* |  | 0.630 | 0.00069 |
| Eco99B829 | 0.27 *a* |  | 0.339 | 0.03689 |
| Pfl29ARP | 0.08 *c* |  | 0.419 | 0.00905 |
| Covariance analysis | df=5, $P=4.4342e-7$,<br>LR=37.65 | df=5, $P=0.83515$,<br>LR=2.1 | | |
| *P+* |  |  |  |  |
| Non-inoculated | 1.15 | 0.60 *c* | 0.399 | 0.00018 |
| AviF0819 | 0.93 | 0.72 *bc* | 0.312 | 0.00132 |
| BveFZB42 | 0.45 | 0.99 *ab* | 0.621 | 2.3175e-7 |
| BveGB03 | 0.79 | 0.81 *bc* | 0.512 | 1.2825e-5 |
| Eco99B829 | 0.17 | 1.15 *a* | 0.312 | 0.00165 |
| Pfl29ARP | 0.10 | 1.15 *a* | 0.529 | 5.3439e-6 |
| Covariance analysis | / | df=5, $P=0.00842$,<br>LR=15.5 | | |

### 3.4 *Brachypodium* total root length and root system morphology were impacted by P supply and inoculation with P solubilizing bacteria

*Brachypodium* TRL increased by 8.97% when plants were exposed to the P-/HA treatment compared to the P+ treatment (*P*=0.023, Fig. 4a-d). In comparison with the TRL measured in non-inoculated plants, the TRL of plants inoculated with BveFZB42, Eco99B829 or Pfl29ARP decreased by 9.64%, 11.61% and 16.67% respectively, whatever the nutritional context (*P*<0.001, Fig. 4e). Mean values per treatment, standard deviation and coefficients of ANOVAs are available in Online Resource 3.

**Fig. 4.**
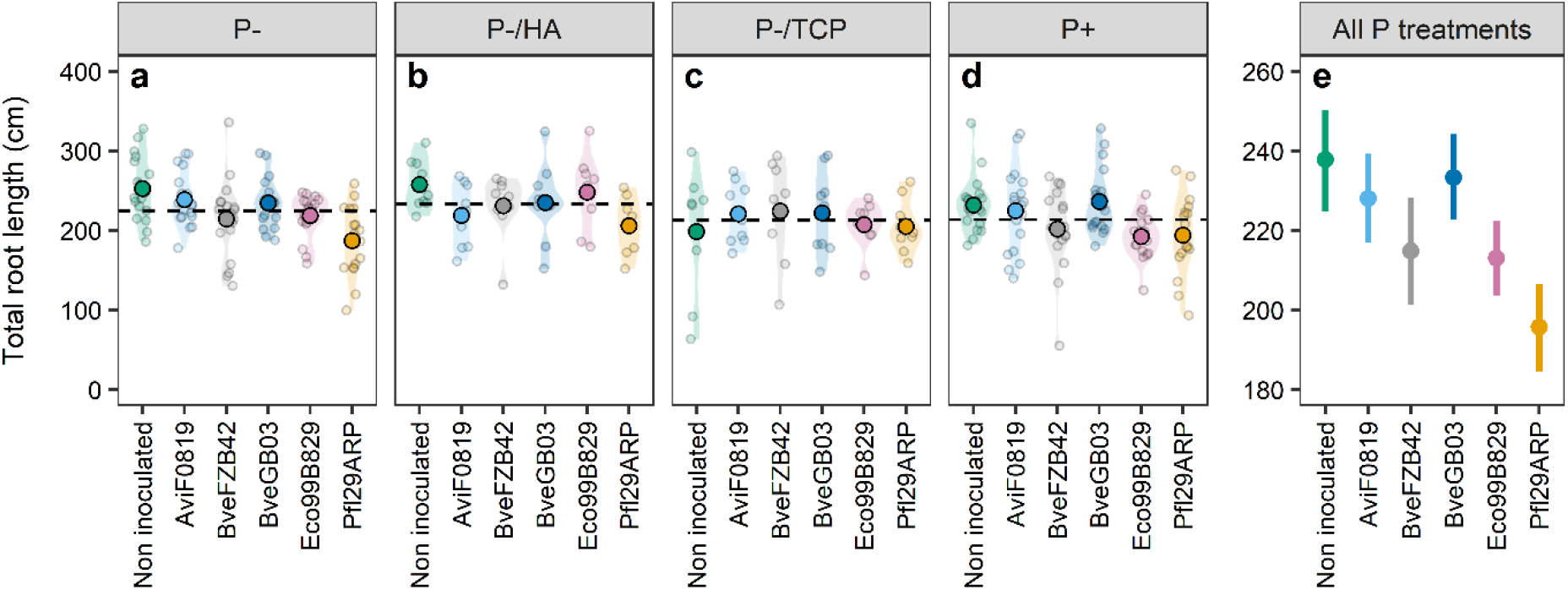
Average total root length of four-week-old *Brachypodium* plantlets exposed to contrasted P supplies and either inoculated or not inoculated with bacteria. n = 18 for the P- and P+ treatments, n = 9 for the P-/HA and P-/TCP treatments. For each P treatment, the grand mean is shown by a dashed horizontal line. For each inoculation treatment, large black-circled dots represent mean values, and shaded areas show the density distribution of each population. Individuals are displayed as small grey-circled dots in the graphs. In panel e, values are means +/- 95% confidence intervals calculated across P treatments

**Online Resource 3.** TRL of four-week-old *Brachypodium* plantlets grown in Magenta boxes, exposed to contrasted P supplies and either inoculated or not inoculated with bacterial strains (n=18 for the P- and P+ treatments, n=9 for the P-/HA and P-/TCP treatments). Results of 3-way ANOVAs (degree of freedom “df”, *P* and *F* values) and Dunnett’s *post hoc* tests (annotated with stars; P+ and non-inoculated treatments used as references)

|  | TRL (cm) |  |
| --- | --- | --- |
|  | mean | sd |
| *P treatment* |  |  |
| P- | 224.58 | 42.78 |
| P-/HA | 233.06* | 42.37 |
| P-/TCP | 213.26 | 49.83 |
| P+ | 213.87 | 46.39 |
| ANOVA | df=3, *P*=0.02126, *F*=3.2821 |  |
| *Inoculation treatment* |  |  |
| Non-inoculated | 237.85 | 48.83 |
| AviF0819 | 228.11 | 42.30 |
| BveFZB42 | 214.92* | 50.62 |
| BveGB03 | 233.41 | 40.92 |
| Eco99B829 | 213.11* | 35.64 |
| Pfl29ARP | 195.82* | 41.59 |
| ANOVA | df=5, *P*=1.99e-06, *F*=7.2513 |  |
| *Repetition* |  |  |
| ANOVA | df=1, *P*=0.07758, *F*=3.1363 |  |
| *Interaction* |  |  |
| P- non-inoculated | 252.83 | 40.23 |
| P- AviF0819 | 239.12 | 33.89 |
| P- BveFZB42 | 214.76 | 49.38 |
| P- BveGB03 | 234.62 | 31.18 |
| P- Eco99B829 | 218.86 | 26.65 |
| P- Pfl29ARP | 187.32 | 42.97 |
| P-/HA non-inoculated | 257.96 | 31.15 |
| P-/HA AviF0819 | 219.12 | 39.79 |
| P-/HA BveFZB42 | 231.37 | 40.44 |
| P-/HA BveGB03 | 235.36 | 49.72 |
| P-/HA Eco99B829 | 248.26 | 45.64 |
| P-/HA Pfl29ARP | 206.27 | 33.97 |
| P-/TCP non-inoculated | 198.82 | 76.65 |
| P-/TCP AviF0819 | 221.07 | 38.39 |
| P-/TCP BveFZB42 | 224.41 | 61.96 |
| P-/TCP BveGB03 | 222.32 | 51.99 |
| P-/TCP Eco99B829 | 207.66 | 28.38 |
| P-/TCP Pfl29ARP | 205.30 | 33.29 |
| P+ non-inoculated | 232.32 | 36.08 |
| P+ AviF0819 | 225.11 | 52.78 |
| P+ BveFZB42 | 202.13 | 51.09 |
| P+ BveGB03 | 236.76 | 41.63 |
| P+ Eco99B829 | 192.52 | 27.13 |
| P+ Pfl29ARP | 194.36 | 47.95 |
| ANOVA | df=15, *P*=0.15092, *F*=1.3891 |  |

The persistent homology analysis of the root systems was performed on 1^st^ and 2^nd^ order roots of non-inoculated plants and plants inoculated with Pfl29ARP or BveFZB42, as those strains showed a strong impact on root biomass accumulation (Fig. 5). Both PSB inoculation and P treatment had a significant impact on root system morphology (*P*<0.001 and = 0.006 respectively). Pairwise comparisons revealed that, on average, the morphology of plant root systems inoculated with Pfl29ARP was different from those of non-inoculated plants and plants inoculated with BveFZB42. Despite a significant impact of P treatment on root system morphology, pairwise comparisons did not highlight the P treatments that differed from one another. The coefficients of the statistical analysis are available in Online Resource 4.

**Fig. 5.**
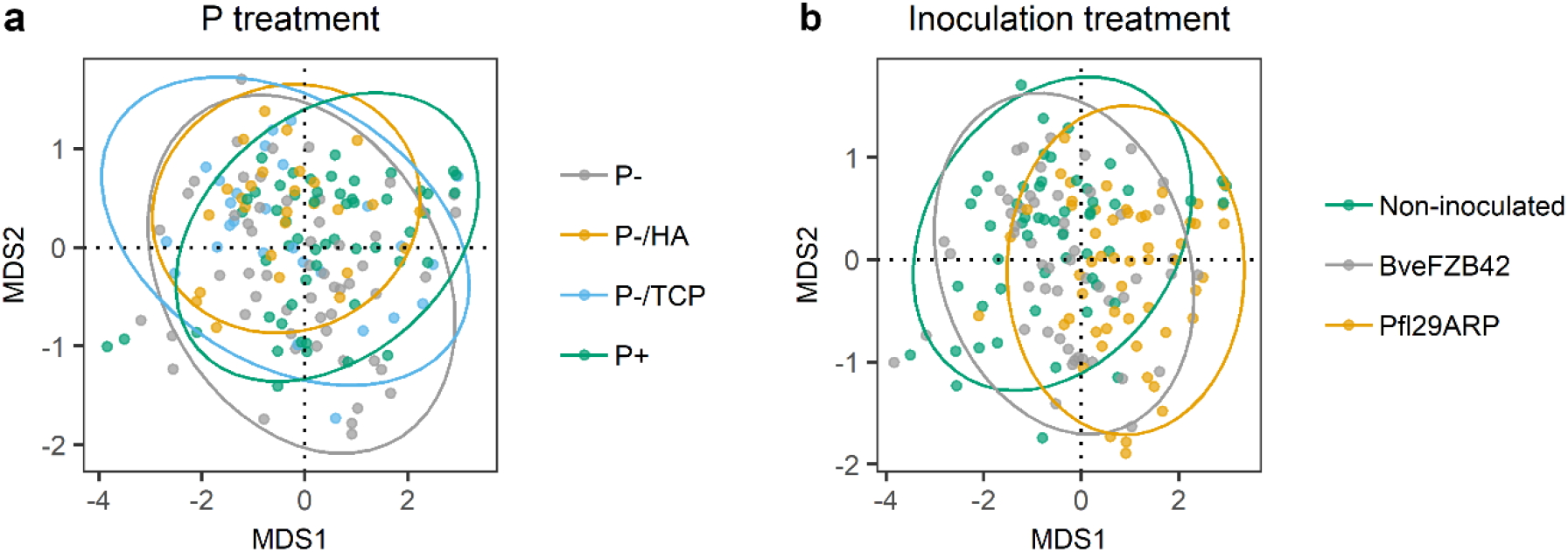
Multidimensional scaling plots displaying morphological differences between root systems, induced by P (a) and inoculation (b) treatments. The Euclidean distance separating two branching structures (dots) on the plot is a close representation of the true dissimilarity between these structures. 95% confidence ellipses for the centroids are plotted for each treatment

**Online Resource 4.** Results of PERMANOVA performed on the persistent homology analysis output of plant root systems. n=18 for the P- and P+ treatments, n=9 for the P-/HA and P-/TCP treatments. Post-hoc tests were performed by running a PERMANOVA for each pairwise comparison and *P* values were adjusted for multiple comparisons using the Benferroni method.

|  | Df | *F* model | *P* |
| --- | --- | --- | --- |
| Inoculation treatment | 2 | 11.1650 | 0.000999 |
| P treatment | 3 | 2.8237 | 0.005994 |
| Interaction | 6 | 1.2461 | 0.217782 |
| Residuals | 150 |  |  |

Post-hoc tests:
|  | *F* model | *P* |
| --- | --- | --- |
| P- vs P-/TCP | 1.8911 | 0.68931 |
| P- vs P+ | 2.1396 | 0.60539 |
| P- vs P-/HA | 2.5906 | 0.28771 |
| P-/TCP vs P+ | 3.1768 | 0.19780 |
| P-/TCP vs P-/HA | 1.4693 | 1.00000 |
| P+ vs P-/HA | 3.5882 | 0.17982 |
| Pfl29ARP vs non-inoculated | 18.7287 | 0.00099 |
| Pfl29ARP vs BveFZB42 | 13.9209 | 0.00099 |
| Non-inoculated vs BveFZB42 | 1.0035 | 0.38462 |

### 3.5 Low P availability induced lower shoot P concentration, even in the presence of P solubilizing bacteria

P concentration in the shoot of plants exposed to the P-, P-/HA and P-/TCP treatments was lower than in plants exposed to P+ (−68.9%, −56.2% and −63.2% respectively; *P*<0.001; Fig. 6a-d). Plants grown under these three treatments showed P deficiency symptoms, such as necrosis starting from the apex of mature leaves (Arvalis, Institut du végétal). Inoculation with bacteria did not help the plants to increase the shoot P concentration, even in the presence of the potentially mobilizable P sources TCP or HA (Fig. 6e). Mean values per treatment, standard deviation and coefficients of ANOVAs are available in Online Resource 5.

**Fig. 6.**
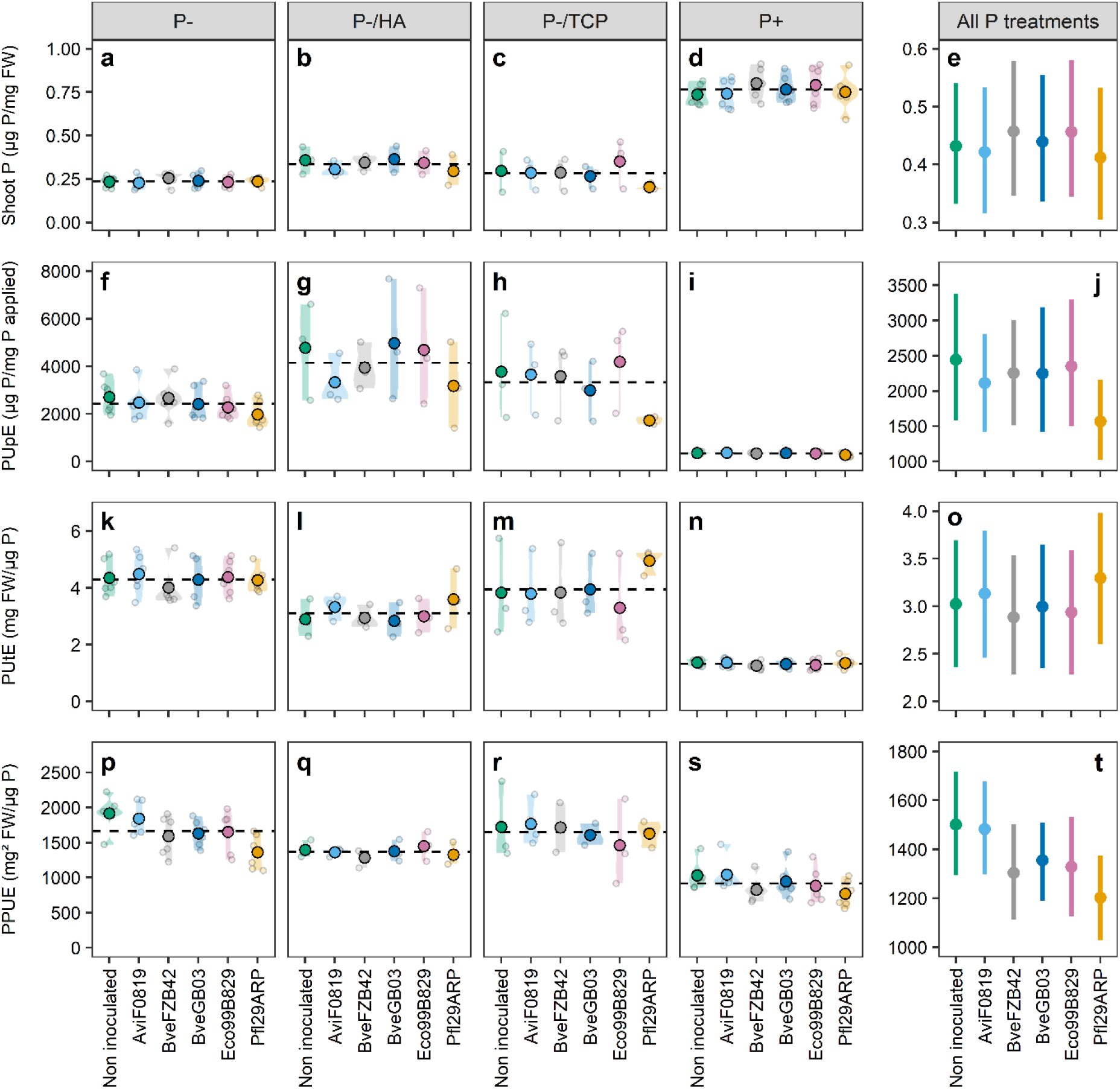
Average shoot P concentration (a-e), P uptake efficiency “PUpE” (f-j), P utilization efficiency “PUtE” (k-o) and physiological P use efficiency “PPUE” (p-t), of four-week-old *Brachypodium* plants grown under contrasted P supplies and either inoculated or not inoculated with bacterial strains. n=6 for the P- and P+ treatments, n=3 for the P-/HA and P-/TCP treatments. For each P treatment, the grand mean is shown by a dashed horizontal line. For each inoculation treatment, large black-circled dots represent mean values, and shaded areas show the density distribution of each population. Individual data points (pool of 5 plantlets) are displayed as small grey-circled dots in the graphs. In panels e, j, o and t, values are means +/- 95% confidence intervals calculated across P treatments

**Online Resource 5.** Shoot P concentration and PUE parameters of four-week-old *Brachypodium* plantlets grown in Magenta boxes, exposed to contrasted P supplies and either inoculated or not inoculated with bacterial strains (n=6 for the P- and P+ treatments, n=3 for the P-/HA and P-/TCP treatments). Results of 3-way ANOVAs (degree of freedom “df”, *P* and *F* values) and Dunnett’s *post hoc* tests (annotated with stars; P+ and non-inoculated treatments used as references)

|  | Shoot P concentration<br>(µg P/mg FW) |  | PU<sub>p</sub>E |  | PU<sub>t</sub>E |  | PPUE |  |
| --- | --- | --- | --- | --- | --- | --- | --- | --- |
|  | mean | sd | mean | sd | Mean | Sd | mean | sd |
| *P treatment* |  |  |  |  |  |  |  |  |
| P- | 0.237* | 0.032 | 2418.692* | 662.318 | 4.296* | 0.603 | 1664.549* | 294.805 |
| P-/HA | 0.335* | 0.064 | 4146.142* | 1768.202 | 3.099* | 0.626 | 1365.267* | 135.433 |
| P-/TCP | 0.281* | 0.092 | 3318.446* | 1565.915 | 3.940* | 1.267 | 1649.241* | 361.100 |
| P+ | 0.764 | 0.086 | 337.427 | 60.022 | 1.325 | 0.151 | 915.617 | 222.696 |
| ANOVA | df=3, *P*<2e-16,<br>*F*=292.0433 |  | df=3, *P*<2e-16,<br>*F*=306.9495 |  | df=3, *P*<2e-16,<br>*F*=217.6723 |  | df=3, *P*<2.2e-16,<br>*F*=60.9819 |  |
| *Inoculation treatment* |  |  |  |  |  |  |  |  |
| Non-inoculated | 0.432 | 0.233 | 2444.236 | 2012.003 | 3.026 | 1.488 | 1501.401 | 467.108 |
| AviF0819 | 0.421 | 0.243 | 2110.551 | 1542.716 | 3.136 | 1.493 | 1482.431 | 419.206 |
| BveFZB42 | 0.457 | 0.260 | 2255.018 | 1662.922 | 2.886 | 1.410 | 1304.184 | 436.268 |
| BveGB03 | 0.440 | 0.248 | 2247.915 | 1942.801 | 2.997 | 1.443 | 1354.915 | 360.533 |
| Eco99B829 | 0.457 | 0.261 | 2350.842 | 2048.284 | 2.937 | 1.485 | 1329.075 | 453.353 |
| Pfl29ARP | 0.412 | 0.256 | 1568.265* | 1258.735 | 3.298 | 1.543 | 1202.833* | 381.678 |
| ANOVA | df=5, *P*=0.2825,<br>*F*=1.2751 |  | df=5, *P*=0.01069,<br>*F*=3.2079 |  | df=5, *P*=0.4357,<br>*F*=0.9788 |  | df=5, *P*=0.007738,<br>*F*=3.3921 |  |
| *Repetition* |  |  |  |  |  |  |  |  |
| ANOVA | df=1, *P*=0.6384,<br>*F*=0.2225 |  | df=1, *P*=0.83696,<br>*F*=0.0426 |  | df=1, *P*=0.4312,<br>*F*=0.6257 |  | df=1, *P*=0.648586,<br>*F*=0.2092 |  |
| *Interaction* |  |  |  |  |  |  |  |  |
| P- non-inoculated | 0.234 | 0.031 | 2704.936 | 682.528 | 4.350 | 0.608 | 1913.742 | 245.850 |
| P- AviF0819 | 0.228 | 0.037 | 2476.670 | 739.706 | 4.487 | 0.686 | 1841.606 | 221.698 |
| P- BveFZB42 | 0.255 | 0.037 | 2664.388 | 746.187 | 4.009 | 0.710 | 1588.721 | 289.616 |
| P- BveGB03 | 0.239 | 0.040 | 2412.522 | 692.725 | 4.284 | 0.701 | 1631.159 | 186.833 |
| P- Eco99B829 | 0.232 | 0.033 | 2275.638 | 540.169 | 4.378 | 0.607 | 1649.714 | 296.266 |
| P- Pfl29ARP | 0.236 | 0.022 | 1977.999 | 544.750 | 4.268 | 0.430 | 1362.351 | 248.987 |
| P-/HA non-inoculated | 0.357 | 0.079 | 4775.731 | 2043.682 | 2.896 | 0.664 | 1395.929 | 126.322 |
| P-/HA AviF0819 | 0.305 | 0.044 | 3327.955 | 1067.802 | 3.323 | 0.456 | 1361.300 | 58.025 |
| P-/HA BveFZB42 | 0.344 | 0.046 | 3942.647 | 989.521 | 2.944 | 0.416 | 1283.964 | 129.002 |
| P-/HA BveGB03 | 0.363 | 0.076 | 4969.660 | 2542.434 | 2.836 | 0.608 | 1374.750 | 154.550 |
| P-/HA Eco99B829 | 0.343 | 0.069 | 4684.583 | 2454.629 | 2.998 | 0.602 | 1449.523 | 212.164 |
| P-/HA Pfl29ARP | 0.295 | 0.089 | 3176.275 | 1807.334 | 3.594 | 1.055 | 1326.138 | 163.137 |
| P-/TCP non-inoculated | 0.296 | 0.117 | 3770.126 | 2229.912 | 3.826 | 1.715 | 1722.268 | 566.133 |
| P-/TCP AviF0819 | 0.285 | 0.089 | 3654.221 | 1536.643 | 3.792 | 1.388 | 1765.498 | 368.753 |
| P-/TCP BveFZB42 | 0.286 | 0.095 | 3586.993 | 1625.492 | 3.835 | 1.530 | 1714.052 | 354.350 |
| P-/TCP BveGB03 | 0.266 | 0.067 | 2989.873 | 1264.261 | 3.944 | 1.112 | 1606.530 | 157.781 |
| P-/TCP Eco99B829 | 0.351 | 0.141 | 4186.080 | 1884.872 | 3.296 | 1.666 | 1460.423 | 611.666 |
| P-/TCP Pfl29ARP | 0.203 | 0.020 | 1723.384 | 157.770 | 4.947 | 0.453 | 1626.672 | 188.967 |
| P+ non-inoculated | 0.736 | 0.062 | 354.844 | 41.169 | 1.368 | 0.114 | 1031.362 | 204.255 |
| P+ AviF0819 | 0.741 | 0.088 | 363.894 | 57.738 | 1.365 | 0.163 | 1042.288 | 220.108 |
| P+ BveFZB42 | 0.802 | 0.086 | 335.845 | 48.824 | 1.260 | 0.138 | 824.824 | 180.071 |
| P+ BveGB03 | 0.765 | 0.078 | 351.455 | 70.629 | 1.318 | 0.129 | 942.947 | 232.530 |
| P+ Eco99B829 | 0.791 | 0.111 | 341.557 | 38.802 | 1.286 | 0.186 | 882.537 | 248.321 |
| P+ Pfl29ARP | 0.751 | 0.102 | 276.967 | 73.901 | 1.354 | 0.193 | 769.744 | 189.371 |
| ANOVA | df=15, *P*=0.5820,<br>*F*=0.8858 |  | df=15, *P*=0.72824,<br>*F*=0.7489 |  | df=15, *P*=0.9355,<br>*F*=0.4976 |  | df=15, *P*=0.937471,<br>*F*=0.4938 |  |

### 3.6 P supply and inoculation with Pfl29ARP impacted P use efficiency components in *Brachypodium*

Regarding the PUpE (i.e. the ratio between shoot P content and applied soluble P), plants exposed to soluble P deficiency had a greater uptake efficiency (*P*<0.001), with the greatest values measured on plants grown in the presence of TCP and HA (883.5% and 1128.8% increase, respectively, compared to plants exposed to P+; Fig. 6f-i). *Brachypodium* acquired and accumulated a greater amount of P in shoots when TCP or HA were added to the nutrient solution, in comparison with the P- treatment and regardless of the bacterial treatment. Plants inoculation with Pfl29ARP led to lower PUpE values under all P treatments compared to non-inoculated plants (average decrease of 35.8% across all P treatments, *P*=0.011, Fig. 6j).

Plants grown under soluble P deficiency were more efficient at utilizing P for biomass accumulation (PUtE, biomass produced by unit of plant P content; *P*<0.001; Fig. 6k-n). These plants accumulated more shoot biomass per unit of shoot P content compared to plants exposed to P+ condition. Plants grown under the P- treatment were globally the most efficient. Inoculation of *Brachypodium* by any of the bacterial strains had no significant impact on PUtE (*P*=0.436, Fig. 6o).

The PPUE (i.e. shoot biomass divided by shoot P concentration), was significantly higher in plants grown under P-, P-/HA and P-/TCP conditions compared to plants exposed to sufficient P supply (81.8%, 49.1% and 80.1% increase respectively compared to P+ condition, *P*<0.001, Fig. 6p-s). Plants exposed to a deficiency in soluble P produced shoot biomass more efficiently at lower shoot P concentration. The inoculation with Pfl29ARP induced a 19.9% reduction in PPUE compared to non-inoculated plants (*P*=0.008, Fig. 6t). Mean values per treatment, standard deviation and coefficients of ANOVAs are available in Online Resource 5.

## 4 Discussion

This study aimed to explore the impact of PSB inoculation on the response of *Brachypodium distachyon* Bd21 to contrasted P conditions. *Brachypodium* and the PSB were co-cultivated over four weeks in an *in vitro* gnotobiotic system and exposed to four different nutritional conditions: a low level of soluble P (P-); a low level of soluble P supplemented with poorly soluble forms of P (P-/TCP and P-/HA); and a high level of soluble P (P+). The plant biomass production and allocation, the root system architecture and the P use efficiency were studied.

### 4.1 *Brachypodium* shows developmental plasticity in response to contrasted P conditions

Our study demonstrated that *Brachypodium* biomass accumulation is highly responsive to P supply, with lower shoot biomass but stable or greater root biomass accumulation under soluble P deficiency compared to high soluble P levels. The reduction in shoot biomass under soluble P deficiency was also reported in *Brachypodium* (Bd21-3) by Poiré et al. (2014) and *Dactylis glomerata* by Haling et al. (2016). Interestingly, a reduction in root biomass was observed in their studies, while our results showed no impact or even an increase in root biomass accumulation when plants were exposed to soluble P deficiency. The younger growth stage obtained in our confined experimental system could explain these results, as the stress was not as intense as it would have been under, for example, greenhouse conditions. Similarly to our results, Giles et al., (2017) showed a decrease in shoot dry weight and an increase in root dry weight in hydroponically grown barley under P deficiency. The stimulation of root development under low P conditions is a common reported response, facilitating the plant to explore the substrate and take up P (Lynch et al., 2012). Nonetheless, it seems that both the growing conditions and the plant growth stage are important factors affecting biomass accumulation in response to P deficiency.

*Brachypodium* displayed different allometric trajectories under contrasted P conditions, showing responsiveness of the allocation pattern to the P supply. Plants grown in the presence of TCP or HA exhibited a higher shoot development per unit of root biomass than plants grown under the P- and P+ treatments. From this we can infer that the presence of unavailable but potentially mobilizable P sources induced a reduction of investment into the root compartment, in comparison with plants grown under P- conditions. Nevertheless, for similar root biomass, the shoot biomass was the highest in plants supplied with the P+ treatment compared to the three other treatments. We can hypothesize that stressed plants (P- conditions) maintained root development at the expense of the shoot compartment. This is confirmed by the greater RMF observed under soluble P limitation. On the contrary, when there was no nutrient limitation, there was no need for the plants to prioritize extension of their root systems and the plants maintained the biomass accumulation into the above-ground compartment. These results are in accordance with the “functional equilibrium model”, which states that a plant shifts allocation towards the organ involved in the acquisition of the most limiting resources (Brouwer 1963), and reveal a true plasticity in response to P supply. Contrasted results were found in previous studies about the allocation pattern in response to P nutrition. Some of them concluded in a “conservative response” of the plants adjusting their size rather than their allocation pattern (apparent plasticity; Müller et al. 2000). Others described an impact on the allocation pattern, but only under severe P stress (Rubio et al. 2013) or in interaction with nitrogen fertilisation (Sims et al. 2012). Plasticity of biomass allocation was also demonstrated, with a strong impact from the nutritional context (Poorter et al. 2012; Poorter and Nagel 2000; Shipley and Meziane 2002).

Regarding the root system, plants exposed to the P-/HA treatment exhibited a greater TRL. The observed root system lengthening was associated with greater root biomass and RMF for plants grown under P- conditions. These results are consistent with those of a hydroponics experiment on several barley varieties, which revealed a general trend towards root lengthening in response to P deficiency (Giles et al. 2017). On the other hand, Shen et al. (2018) reported that under moderate P stress, wheat plants maintained root length and reduced root biomass whereas under severe P stress both TRL and root biomass were reduced.

### 4.2 Despite their ability to solubilize tricalcium phosphate and hydroxyapatite, the bacterial inoculants did not alleviate P deficiency stress in *Brachypodium* under the experimental growing conditions

All the selected bacteria were able to solubilize the poorly available forms of P (TCP and HA) in NBRIP medium (Nautiyal 1999). HA, despite being reported as less soluble than TCP (Bashan et al. 2013, Havlin et al. 2014), was as easily solubilized as TCP. Some acidification of the medium was observed, with the best solubilizer strains acidifying the most. Medium acidification by proton release is the most straightforward P solubilization process (Bashan et al. 2013) and numerous studies have reported an acidification-associated P solubilization (Collavino et al. 2010; Fernández et al. 2012; Pereira and Castro 2014; Yu et al. 2011). The relationship between soluble P concentration and pH variation tended towards an intensification of P solubilization activity as the pH variation became stronger. This was more pronounced for HA than for TCP solubilization. This raises the hypothesis that HA solubilization mechanisms other than acidification are involved, such as complexing or chelating reactions (Bashan et al. 2013).

The use of PSB as bio-inoculants is increasingly reported in the literature, with interesting effects of microbial P mobilization on plant development and yield (Bakhshandeh et al. 2015; Li et al. 2017; Oteino et al. 2015; Pereira and Castro 2014), but few results have reported the inefficiency of *in vitro*-selected PSB to promote plant growth in the presence of poorly soluble forms of P (Collavino et al. 2010; Yu et al. 2011). In our study, the biomass accumulated in shoots and roots was reduced when plants were grown in the presence of bacteria. The strains Pfl29ARP and BveFZB42 had the strongest impact on plant development. Despite their ability to solubilize TCP and HA in NBRIP medium, the selected strains were not able to mobilize these poorly soluble forms of P under co-cultivation conditions and by this way alleviate P-starvation stress in *Brachypodium*. The soluble P concentration in the Hoagland solution at the end of the cultivation was below the detection limit of our analytical method for the P-, P-/TCP and P-/HA treatments (data not shown). A slight acidification of the nutrient solution was observed at the end of the co-cultivation in the presence of bacteria, but the pH remained within an acceptable range for plant development (data not shown). The available carbon source is of great importance for P solubilization by bacteria. Nico *et al*. (2012) reported a reduced P solubilization by bacteria unless glucose was added to the growing medium of rice. Soil experiments also resulted in the absence of beneficial effects of microbial consortia products on maize grown in a substrate with low organic matter content (Bradácová et al. 2019). Glucose is the most abundant sugar detected in *Brachypodium* exudates (Kawasaki et al. 2016). In our study, the bacterial strains were tested for P solubilization with glucose as the sole C source in NBRIP medium (Nautiyal 1999). During the co-cultivation experiment, the concentration of glucose provided through root exudates may have been too low to sustain the bacteria solubilization activity. As our gnotobiotic co-cultivation system was closed during the entire experiment duration, toxic bacterial metabolites may have accumulated in the system, leading to a repression of plant growth. Some studies have revealed a deleterious impact of inoculation with bacterial strains on plant growth under gnotobiotic conditions (Rybakova et al. 2016; Timmusk et al. 2015). The efficacy of a system to test for PSB activity in the presence of a host plant appears to be highly dependent on the considered organisms, but also on the co-cultivation conditions.

### 4.3 The plastic response of *Brachypodium* to P deficiency was modulated by inoculation with P solubilizing bacteria

Regarding the biomass allocation pattern, inoculation with PSB revealed an alteration of the plant’s response to P conditions, except in the presence of HA. Under P- conditions, inoculation with PSB (except with BveFZB42) led to a reduced shoot productivity for similar root biomass. The same observation was made under the P-/TCP treatment, mainly with Pfl29ARP. The depletion in shoot growth benefited the root system, the development of which was either unaffected or less impacted than the shoot. This resulted in an increase in RMF. Under the P+ treatment, investment into the root compartment was reduced in inoculated plants, except with AviF0819 and BveGB03. The RMF was still increased for the same reason as before: a repression of shoot biomass but a steady root biomass accumulation. As the root system is the place where the interaction with the bacteria occurs, it appears that the plant modulated the development of this interface of interaction depending on the nutritional context. These contrasted behaviours in *Brachypodium* should be explored more deeply. The complementarity between biomass partitioning (RMF) and allometric trajectories appears clearly here for the analysis of biomass allocation patterns under environmental variation. Both approaches should be considered when studying the impact of biostimulants on plant biomass allocation in response to environmental constraints.

The total root length of *Brachypodium* was significantly impacted by the P supply and inoculation with PSB. Regardless the P treatment, inoculation with BveFZB42, Eco99B829 and Pfl29ARP led to a reduction in TRL. These results contrast with others reported in the literature. Indeed, Talboys et al. (2014) demonstrated a root elongation promotion effect of BveFZB42 inoculation on wheat (through auxin production), in both low and high P-level soils. In a soil experiment, *Pseudomonas fluorescens* strains also exhibited a positive impact on wheat root elongation under contrasted P fertilisation (Zabihi et al. 2011). The persistent homology analysis performed in our study revealed that inoculation with Pfl29ARP impacted the morphology of the plant root system (considering 1^st^ and 2^nd^ order roots) in comparison with non-inoculated plants and plants inoculated with BveFZB42. The P conditions also induced changes in root system morphology, but these were less easily characterized. According to our results, *Brachypodium* showed a modification of root development, triggered by contrasted P supply and inoculation with bacteria. Geometrical and topological aspects of the root system architecture are important for nutrient foraging in soils. Both aspects are covered by the persistent homology analysis of the root system morphology. Thus, the methodological approaches used in this study appear suitable in seeking to characterize the plant’s response to P supply and inoculation with PSB. This study did not consider root hairs, yet they constitute an important strategy for P nutrition (Lynch 2011) and should be further investigated.

### 4.4 Inoculation with P solubilizing bacteria did not improve *Brachypodium* P use efficiency under the experimental growth conditions

The shoot P concentration and PUE in *Brachypodium* were mainly affected by the P supply, but also by PSB inoculation to some extent. The shoot P concentration was the lowest in plants grown under P- conditions, confirming the P-deficient status of those plants. Despite the demonstrated ability of the bacterial strains to solubilize TCP and HA, they did not alleviate P deficiency in the plants. The soluble P concentration in the Hoagland solution at the end of the cultivation was null for the P-, P-/TCP and P-/HA treatments (data not shown). This result reinforces the above-mentioned hypothesis that the PSB did not extensively solubilize TCP and HA in our gnotobiotic conditions. On the other hand, the P+ solution contained enough soluble P after four weeks for avoiding nutritional stress in the plants (data not shown). Considering the slightly higher shoot P concentration in the presence of TCP and HA regardless the inoculation treatment, we assume that *Brachypodium* was able to partly solubilize those poorly soluble forms of P. Indeed, plants are able to acidify the rhizosphere and release organic anions, mobilizing poorly available P sources (Hinsinger et al. 2003; Wang and Lambers 2019). Citrate, malate, succinate, fumarate and oxalate were detected in *Brachypodium* root exudates (Kawasaki et al. 2016) and may be implied in P solubilization processes. The PUpE was significantly higher in plants exposed to soluble P deficiency compared to plants grown under the P+ treatment, as the stressed plants took up all the available soluble P and partly used it to build their shoots. The highest PUpE values were obtained in the presence of TCP and HA. This observation is consistent with the higher shoot P concentration observed under these treatments and reinforces the above-mentioned hypothesis of partial P solubilization by *Brachypodium*. The PUpE reduction in plants inoculated with Pfl29ARP is consistent with the observed decrease in shoot biomass accumulation, which impairs their P accumulation ability. The PUtE was significantly higher under soluble P deficiency than under the P+ treatment, with the highest efficiency under the P- treatment. Therefore, stressed plants produced the largest biomass per unit of accumulated P. The inoculation of *Brachypodium* with bacteria did not impact the PUtE, as expected from their poor P solubilization activity during the co-cultivation experiment. As observed for PUpE and PUtE, the PPUE values were higher under soluble P deficiency, meaning that for similar shoot P concentration the stressed plants produced more shoot biomass. The inoculation with Pfl29ARP induced a reduction in PPUE. Indeed, shoot P concentration was similar in non-inoculated plants and in plants inoculated with Pfl29ARP, but shoot biomass accumulation was reduced in inoculated plants.

## 5 Conclusion

The selected PSB efficiently solubilized TCP and HA in an *in vitro* liquid cultivation system. However, they did not alleviate P deficiency in *Brachypodium* under gnotobiotic co-cultivation conditions. Some negative impact of the PSB on plant biomass accumulation was even observed, probably due to inadequate carbon supply through root exudates or to the accumulation of bacterial toxic metabolites in the system. *Brachypodium* showed developmental plasticity in response to contrasted P conditions, prioritizing the development of the root compartment upon P starvation. Despite their inability to alleviate P deficiency, the selected PSB modulated *Brachypodium*’s response to P conditions by altering the plant allocation pattern and the root system development. Nevertheless, this modulation did not improve PUE in *Brachypodium* under our experimental conditions. This study highlights the necessity to select experimental conditions as close as possible to realistic conditions in the perspective of screening PSB for the purpose of using them as plant inoculants. Co-cultivation experiments are mandatory in order to confirm a beneficial interaction and test the related hypothesis. To our knowledge, this study represents the first time that allometry and persistent homology analyses were used to assess the impact of biostimulants on plant development under nutritional deficiency. They revealed to be convenient tools to study potential plasticity in biomass allocation or change in root system morphology. The plasticity in biomass allocation could be explored more deeply by considering a temporal perspective of the biomass allocation patterns; this would allow the experiment to cover a broader range of plant sizes and clearly assess the interaction between the use of biostimulants and varying nutrient supply. As root hairs are an important trait in nutrient acquisition, they deserve consideration in addition to root system architecture parameters, providing a more precise insight into root system plasticity in response to P supply and PSB inoculation. Integrating the proposed analyses and tools in future research would provide a better understanding of the impact of biostimulants on plant plasticity in a changing environment.

## Abbreviations

HA: hydroxyapatite
PPUE: physiological P use efficiency
PSB: phosphate-solubilizing bacteria
PUpE: P uptake efficiency
PUtE: P utilization efficiency
PUE: P use efficiency
RMF: root mass fraction
SMA: standardized major axis
TCP: tricalcium phosphate
TRL: total root length

## 6 Acknowledgments

This research was supported by internal research funds of the University of Liège (Belgium). The authors are thankful to Florence Paquet for her technical support, Dr Yves Brostaux (Gembloux Agro-Bio Tech) for his constructive advice on statistical analyses and Guillaume Lobet (Forschungszentrum Juelich, Germany) for reviewing the manuscript.

